# Primary cilia promote EMT-induced triple-negative breast tumor heterogeneity and resistance to therapy

**DOI:** 10.1101/2024.09.10.612175

**Authors:** Camille E. Tessier, Jennifer Derrien, Aurore M.M. Dupuy, Thomas Pelé, Martin Moquet, Julie Roul, Elise Douillard, Camille El Harrif, Xavier Pinson, Matthieu Le Gallo, Florence Godey, Patrick Tas, Roselyne Viel, Claude Prigent, Éric Letouzé, Peggy Suzanne, Patrick Dallemagne, Mario Campone, Robert A. Weinberg, Jacqueline A. Lees, Philippe P. Juin, Vincent J. Guen

## Abstract

Tumor heterogeneity and plasticity, driven by Epithelial-Mesenchymal Transition (EMT), enable cancer therapeutic resistance. We previously showed that EMT promotes primary cilia formation, which enables stemness and tumorigenesis in triple-negative breast cancer (TNBC). Here, we establish a role for primary cilia in human TNBC chemotherapeutic resistance. We developed patient-derived organoids, and showed that these recapitulated the cellular heterogeneity of TNBC biopsies. Notably, one of the identified cell states bore a quasi-mesenchymal phenotype, primary cilia, and stemness signatures. We treated our TNBC organoids with chemotherapeutics and observed partial killing. The surviving cells with organoid-reconstituting capacity showed selective enrichment for the quasi-mesenchymal ciliated cell subpopulation. Genomic analyses argue that this enrichment reflects a combination of pre-existing cells and ones that arose through drug-induced cellular plasticity. We developed a family of small-molecule inhibitors of ciliogenesis and show that these, or genetic ablation of primary cilia, suppress chemoresistance. We conclude that primary cilia help TNBC to evade chemotherapy.

**Significance:** Cancer cells that activate EMT to acquire a quasi-mesenchymal state form primary cilia to evade chemotherapy in human triple-negative breast cancer. Pharmacological inhibition of primary ciliogenesis counteracts EMT-induced chemoresistance.

## Introduction

Tumor heterogeneity and plasticity contribute to cancer therapeutic resistance which is responsible for the death of most cancer patients (*1–4*). Resistance of cancer cells to treatment can result from both primary and acquired resistance to therapy (*2, 4–7*). Epithelial-Mesenchymal Transition (EMT) is a cell biological program that drives tumor heterogeneity (*2, 8, 9*), enabling epithelial cancer cells to acquire an array of mesenchymal phenotypes (*10–12*). Tumor cells that activate this cell plasticity program can transit from early to intermediate to late hybrid epithelial-mesenchymal (E/M) states before reaching a fully elongated mesenchymal morphology, an endpoint that is rarely reached in spontaneously arising tumors (*8, 10*). The distinct phenotypic states arrayed along the E-to-M axis are referred to as EMT states and cells in each of these states can display distinct functional properties (*10, 13, 14*). Cells residing in the late hybrid E/M state, also known as the quasi-mesenchymal state, display tumor-initiating, metastatic, and therapy-resistant properties and are thought to function as cancer stem cells in many carcinomas (*8, 9, 15*).

The degree to which EMT is activated in the neoplastic cells of a tumor is thought to rely on its cell-of-origin, which influences tumor heterogeneity and impacts response to therapy (*16, 17*). Pioneering work revealed that the EMT program endows carcinoma cells with chemotherapy resistance properties, doing so by influencing the expression of ABC drug efflux pumps (*18, 19*), antioxidant enzymes (*19*), pro-apoptotic proteins (*20*), or by promoting DNA damage repair (*21*). Other important studies revealed that EMT can also trigger cancer cell resistance to targeted therapies, by promoting a shift in cell signaling dependencies (*22, 23*), as well as to immunotherapy (*24*), the latter achieved by orchestrating changes in tumor immune microenvironment (*25*). Despite progress in understanding EMT-induced therapeutic resistance with the help of mouse tumor models and cancer cell lines *in vitro* (*26*), there is still limited understanding of the degree to which EMT programs contribute to inter- and intratumor phenotypic heterogeneity during human tumor development and responses to therapy.

Our own research has shown that EMT programs are activated in human triple-negative breast cancers (TNBCs) (*27*). We found that EMT programs promote TNBC/claudin-low tumorigenesis by inducing primary ciliogenesis and ciliary signaling in tumor-initiating carcinoma cells (*27, 28*). The primary cilium is a microtubule-based structure which can be assembled as a solitary structure at the surface of cells composing both normal and neoplastic tissues where it serves as a cell signaling hub (*29, 30*). However, the role of primary cilia in cancer pathogenesis and therapeutic response remains poorly understood.

Here, we used patient tumor biopsies and patient-derived cancer organoids (PDOs) to investigate the degree to which the EMT and primary cilia contribute to human TNBC heterogeneity and therapeutic response. TNBCs represent the most aggressive group of human breast tumors. Most of the highly progressed TNBCs display long-term resistance to therapy and therefore represent an unmet medical need despite recent progress in developing effective therapeutic agents (*31*). Research using mouse cancer models and cell lines *in vitro* has suggested that TNBC heterogeneity, tumorigenesis, and therapeutic resistance is driven by EMT, notably in the claudin-low molecular subtype of this disease (*9, 17, 19, 23, 25, 27, 32–38*). In the present work we show that primary cilia promote EMT-induced tumor heterogeneity and resistance to chemotherapy in TNBCs. EMT induces primary ciliogenesis in malignant cells that reside in a quasi-mesenchymal state. Moreover, primary ciliogenesis represents a vulnerability of these therapy-resistant carcinoma cells that are often the source of therapeutic failure.

## Results

### Late hybrid quasi-mesenchymal cancer cells assemble primary cilia in human TNBCs

To investigate the degree of phenotypic heterogeneity of human TNBCs associated with EMT features and primary ciliogenesis, we first used patient-derived tumor biopsies (PDBs). We collected PDBs from 13 patients who had not received therapy at the time of the biopsy. We identified cancerous lesions in PDBs through hematoxylin-eosin staining of sections of tumor biopsies (Supplementary Fig. 1A). In addition, we co-stained adjacent tumor sections for the epithelial marker E-cadherin (Ecad), the mesenchymal marker Vimentin (Vim), and the primary cilium marker ADP-ribosylation factor-like protein 13b (Arl13b, Fig. 1A). We found double-positive Ecad (Ecad+)/Vimentin (Vim+) cancer cells residing in hybrid epithelial-mesenchymal (E/M) states in all samples (Fig. 1B). Importantly, the representation of Ecad+/Vim+ cancer cells varied significantly between patient’s tumors and even between distinct areas within an individual tumor, ranging from 0 to 100 % of Ecad+/Vim+ cancer cells (Fig. 1B). Using the Arl13b staining, we detected primary cilia in 12 of the 13 tumors initially analyzed and at various frequencies in distinct regions of interest within individual tumors and between tumors, ranging from 0 to 17.39 % of cells that were stained (Fig. 1A, C). Most importantly, we found that double-positive Ecad+/Vim+ cancer cells are significantly more ciliated that Ecad+/Vim-cells indicating that the propensity of cancer cells to form primary cilia reflects acquisition of a hybrid E/M state (Fig. 1D).

**Figure 1.**
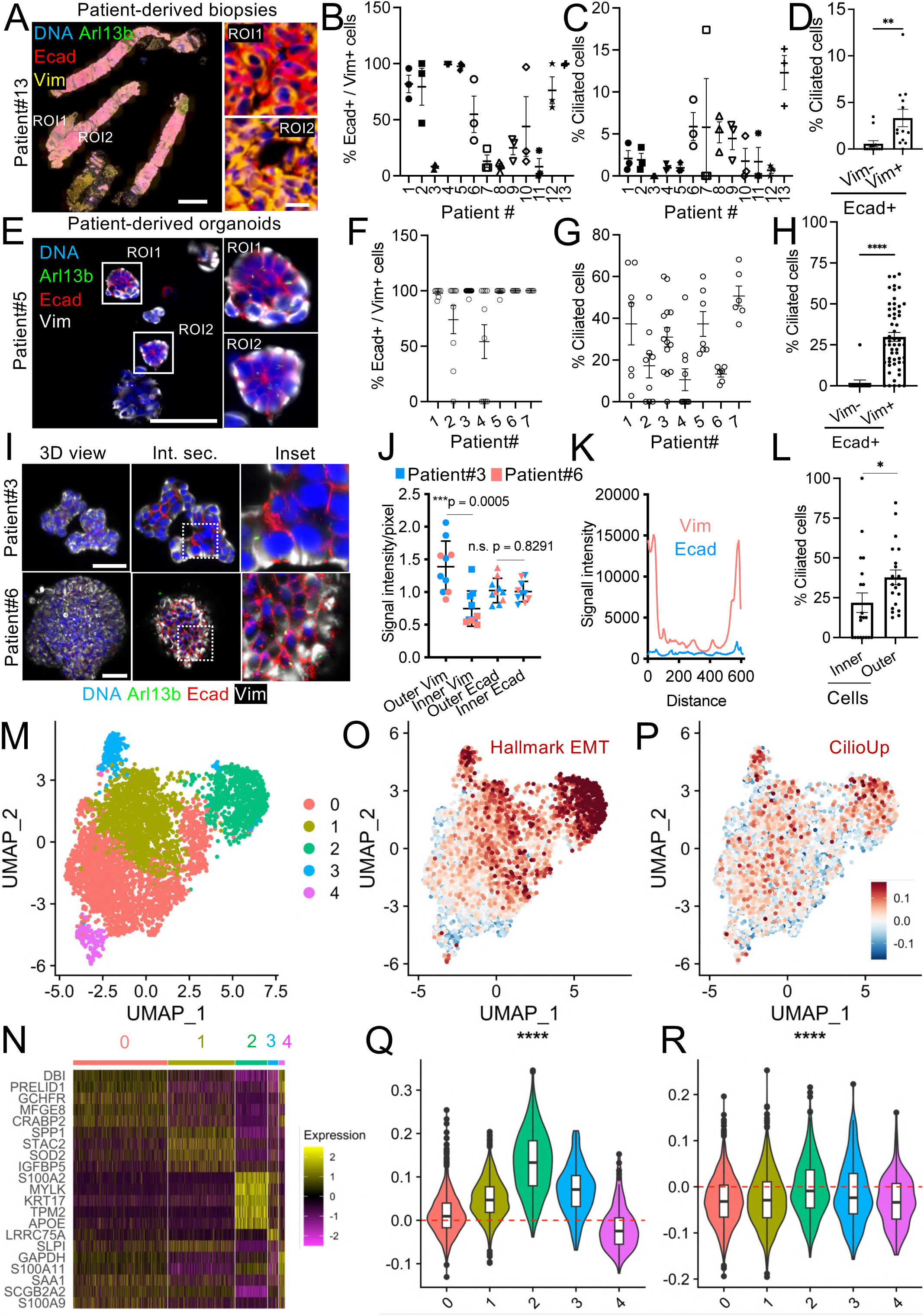
Late hybrid quasi-mesenchymal cancer cells assemble primary cilia in human TNBCs. **(A)** Patient-derived tumor biopsies (PDB) were stained for the indicated proteins (n = 13, representative result for patient#13 is shown). Scale bar: 2 mm (insets: 3x). **(B)** The percentage of Ecad+/Vim+ cells was quantified in three distinct Ecad+ tumor region of interests (ROI) for each PDB (mean ± sem). **(C)** The percentage of Ecad+ ciliated cells was quantified in the same ROIs (mean ± sem). **(D)** The percentage of Ecad+/Vim+ vs Ecad+/Vim-ciliated cells was determined. (n= 13, mean ± sem, **P ≤ 0.01. **(E)** Patient-derived tumor organoids (PDOs) from distinct patient samples were stained for the indicated proteins (n = 7, representative result for patient #5). Scale bar: 100 μm (insets: 3x). **(F-G)** The percentage of Ecad+/Vim+ cells and the percentage of ciliated cells was quantified in optical sections per PDO in each patient sample (n ≥ 5 PDOs/patient sample, mean ± sem). **(H)** The percentage of Ecad+/Vim+ vs Ecad+/Vim-ciliated cells was determined. (n= 68 PDOs, 6 distinct patient samples, mean ± sem, **P ≤ 0.01). **(I)** Whole mount immunofluorescence staining was conducted on PDOs for the indicated markers. 3-dimensional imaging of PDOs was performed. The 3D representation (3D view) as well as internal optical sections (int. sec.) are shown for representative PDOs for two distinct patient samples. Scale bars: 50 μm (inset: 3x). **(J)** Signal intensity for Ecad and Vim staining was measured across cells in individual PDOs for the two distinct patient samples, in cells residing within PDOs (inner) in comparison to cells residing at the periphery of PDOs (outer, n ≥ 5 PDOs/patient sample, mean ± sem). **(K)** Distribution of the Ecad and Vim signal across one representative PDO is shown. **(L)** Ciliation of cells residing within PDOs in comparison to cells on the outer part of PDOs was quantified (n = 20 PDOs, 4 distinct patient samples, mean ± sem; p ≤ 0.05). **(M)** Gene expression in cancer cells of PDOs was analyzed by scRNAseq (n= 4888 cells). UMAP illustrating transcriptional heterogeneity between cancer cells of PDOs. Each point represents a cell colored according to its cell cluster (0–4). **(N)** Heatmap highlighting marker genes of each cluster. **(O-P)** UMAP and violin plots illustrating the expression of EMT and ciliogenesis signatures in the distinct cell clusters.

We next assessed phenotypic heterogeneity linked to EMT and primary ciliogenesis in patient-derived organoids (PDOs) using 7 distinct patient samples (Supplementary Fig. 1B). To begin, we assessed inter- and intraorganoid phenotypic heterogeneity by conducting whole mount immunofluorescence co-staining and imaging of PDOs (Fig. 1E). We detected PDOs that were comprised of double-positive Ecad+/Vim+ cells in all breast tumor-derived samples (Fig. 1F). The representation of Ecad+/Vim+ double-positive cancer cells varied significantly between tumor samples and between PDOs for each tumor sample, ranging from no detectable Ecad+/Vim+ double-positive cells in some PDOs to PDOs exclusively composed of Ecad+/Vim+ cells (Fig. 1F). Of note, we detected more Ecad+/Vim+ cells in PDOs in comparison to PDBs (Fig. 1B, F). We next assessed primary ciliogenesis in PDOs using Arl13b staining (Fig. 1G). Here, we found primary cilia in all breast tumor-derived samples but at different frequencies between PDOs (Fig. 1G). Most importantly, we found that cancer cells that co-expressed Ecad and Vim were significantly more ciliated than cancer cells that expressed exclusively Ecad (Fig. 1H). This revealed that the propensity of cancer cells to form primary cilia follows acquisition of a hybrid E/M phenotype in PDOs, reflecting the earlier findings in PDBs. Together, our findings in PDBs and PDOs establish that inter- and intratumor phenotypic heterogeneity in human TNBCs is linked to the epithelial-mesenchymal status of cancer cells and primary ciliogenesis. Malignant cells that reside in a hybrid E/M state have the ability to form primary cilia. However, acquisition of a hybrid E/M phenotype is not sufficient for primary ciliogenesis and only a subset of hybrid E/M cells form primary cilia.

To gain additional insight into the identity of hybrid E/M cells that form primary cilia, we conducted additional studies using PDOs. To begin, using whole-mount immunofluorescent staining and 3D-imaging studies of these structures, we found that Ecad+ cells that express high levels of Vim that reside on the outer layers of PDOs exhibited an elevated ability to form primary cilia when compared with the inner cells (Fig. 1I-L, Supplementary Movie 1). We next conducted single-cell RNA sequencing (scRNAseq) analysis of cancer cells dissociated from PDOs. Unsupervised clustering of cancer cells revealed five different cell states in the cells forming the analyzed PDOs (Cluster 0-4, Fig. 1M-N, Supplementary Table 1). Remarkably, we found evidence for candidate genetic alterations scattered in cancer cells by inferring copy number alterations (inferCNV) from our scRNA-seq data (Supplementary Fig. 1C). Each copy-number subclone displayed distinct phenotypes corresponding to the distinct cell states previously defined, with varying proportions according to the subclone, indicating that epigenetic events contribute to transcriptional heterogeneity between cancer cells of our PDOs (Supplementary Fig. 1C). To determine whether EMT and ciliogenesis programs contribute to transcriptional heterogeneity of cancer cells, we next assessed expression of transcriptional signatures that mark EMT and ciliogenesis in the distinct cell states (Fig. 1O-R, Supplementary Table 2). Importantly, we found differential expression of the signatures between cancer cell clusters and identified one cell cluster composed of cells that co-expressed the highest level of EMT and ciliogenesis transcriptional signatures (cluster 2, Fig. 1O-R). Additional supervised and unsupervised gene set enrichment analysis revealed other significantly enriched gene sets in the list of genes that are up-regulated in cancer cells of cluster 2, including stem cell transcriptional signatures (Supplementary Fig. 1D-E, Supplementary Table 3). Altogether, our data indicate that cancer cells that reside in a late hybrid quasi-mesenchymal stem-like state represent a subpopulation of hybrid E/M cells that assemble primary cilia in human TNBCs.

### Late hybrid quasi-mesenchymal ciliated cells display enhanced resistance to chemotherapy

To investigate the response of cancer cells residing in distinct states to chemotherapy, we next treated our PDOs with doxorubicin and taxol, two drugs that are in use in the oncology clinic (Supplementary Fig. 2A). We found that both chemotherapeutics induced a reduction in cancer cell viability in a dose-dependent manner (Supplementary Fig. 2A). Most importantly, a subset of cancer cells displayed enhanced chemoresistance at intermediate doses and organoid-reconstituting capacity (Supplementary Fig. 2A-B). Using immunostaining, we found that PDOs comprised of chemoresistant cells are enriched for hybrid E/M cells that assemble primary cilia in comparison with PDOs comprised of vehicle-treated control cancer cells (Fig. 2A). To gain additional insights into the identity and origin of these cells, we next conducted scRNAseq analysis of cancer cells dissociated from vehicle-treated PDOs (CTL) or PDOs post-chemotherapy. Unsupervised clustering of cells revealed the existence of five different cell states in these PDOs (clusters 0-4, Fig. 2B-D, Supplementary Table 4). The percentage of cells from cluster 0 was similar in the control PDOs and PDOs post-chemotherapy (Fig. 2E). However, the representation of cells from cluster 1 and 3 decreased and the representation of cells from cluster 2 and 4 increased in PDOs post-chemotherapy when compared with control PDOs (Fig. 2E). Importantly, cells of cluster 2 and 4 expressed the highest levels of EMT, ciliogenesis, and stem cell transcriptional programs (Fig. 2F-G, Supplementary Fig. 2C-D, Supplementary Table 5). These findings revealed selective enrichment for late hybrid quasi-mesenchymal stem-like ciliated cell subpopulation after chemotherapy.

**Figure 2.**
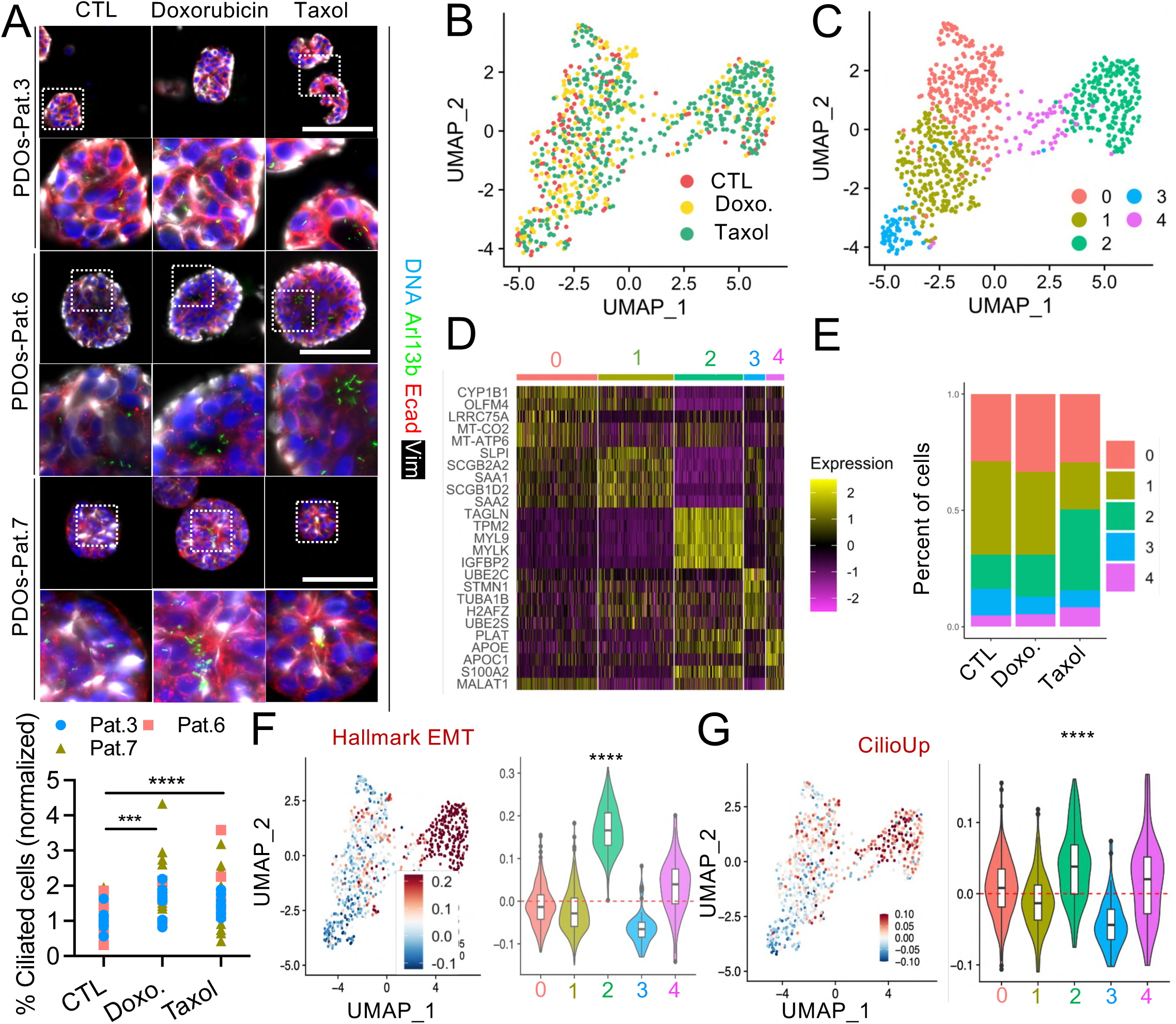
Late hybrid quasi-mesenchymal ciliated cells display enhanced resistance to chemotherapy. **(A)** PDOs from 3 distinct patient (pat.) samples were stained for the indicated proteins. Scale bars: 100 μm (insets: 3x). The percentage of ciliated cells was quantified in optical sections in PDOs for each patient sample (n ≥ 33 PDOs/treatment condition from 3 distinct patient samples, mean ± sem). **(B)** Gene expression in cancer cells of PDOs was analyzed by scRNAseq (n = 820 cells). UMAP plot integrating data from 3 samples (CTL = DMSO treated, Doxo. = doxorubicin, Taxol). Each point represents a cell colored by sample of origin. **(C)** UMAP illustrating the different cell clusters identified by transcriptional heterogeneity. Each point represents a cell colored according to its cell cluster (0–4). **(D)** Heatmap highlighting marker genes of each cluster. **(E)** Percentage of cells in each cluster per sample. **(F-G)** UMAP and violin plots illustrating the expression of EMT and ciliogenesis signatures in the distinct cell clusters.

### Late hybrid quasi-mesenchymal cells arise through drug-induced cellular plasticity associated with activation of EMT and ciliogenesis transcriptional programs

Using inferCNV, we found evidence that at least a subset of cells residing in late hybrid quasi-mesenchymal state after chemotherapy arose through drug-induced cellular plasticity (Fig. 3A-B, S3A-C). Late hybrid quasi-mesenchymal cells of cluster 2 and 4 enriched post-treatment were composed of a subset of cells devoid of a candidate amplification of chr11:45909669-65044828 and of another subset of cells harboring chr11:45909669-65044828 amplification that was only found in more early hybrid E/M cells of cluster 0-1-3 before treatment (Fig. 3A-B, S3A-C). Using pseudotime analysis, we identified a lineage trajectory from cells residing in the early hybrid states (clusters 3-1-0) to cells residing in the late-hybrid state (4–2) associated with progressive expression of EMT and ciliogenesis transcriptional programs (Fig. 3C-F). These data reveal that at least a subset of late hybrid quasi-mesenchymal ciliated cells with enhanced resistance to chemotherapy arose through drug-induced cellular plasticity associated with activation of EMT and primary ciliogenesis transcriptional programs.

**Figure 3.**
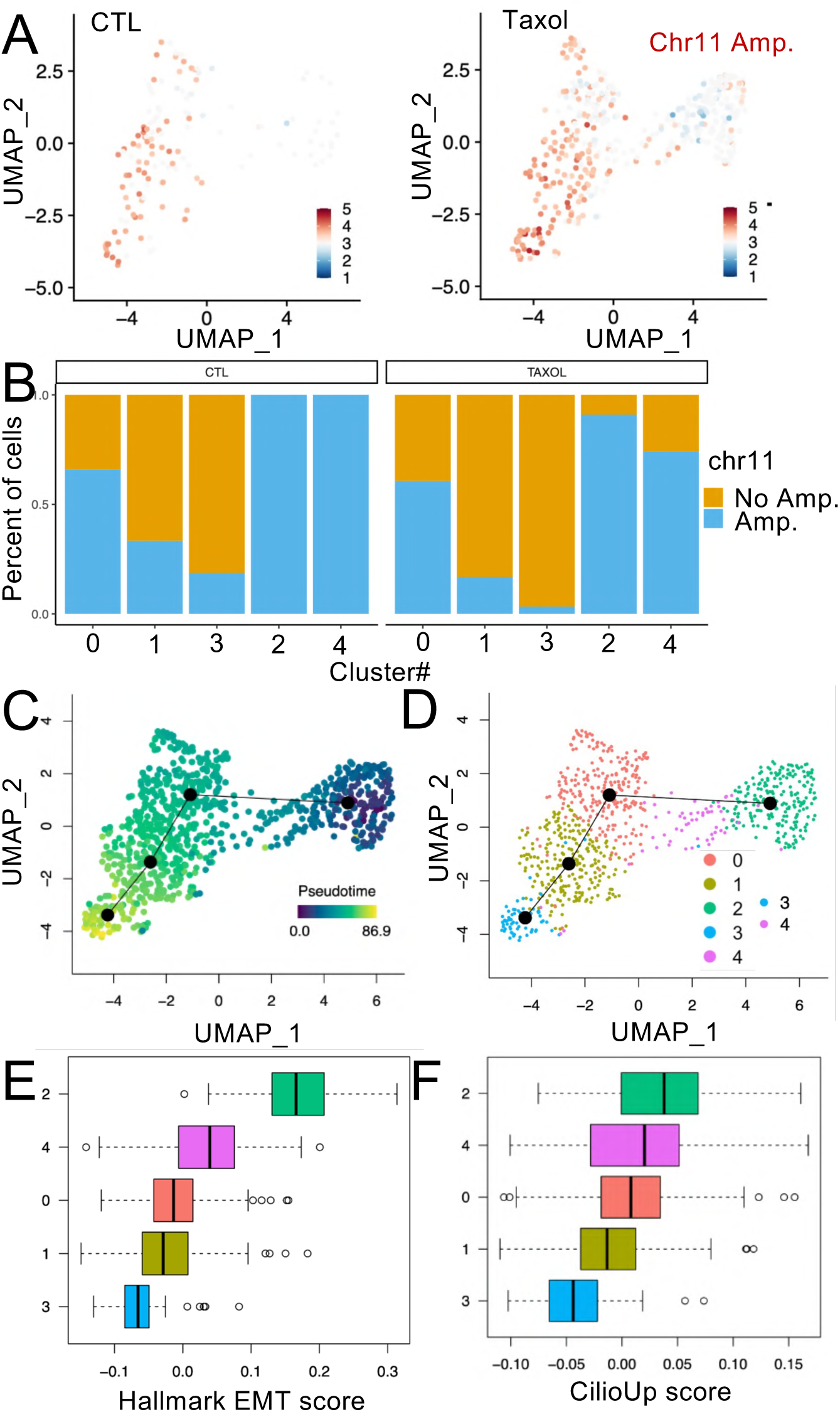
Late hybrid quasi-mesenchymal cells arise through drug-induced cellular plasticity associated with activation of EMT and ciliogenesis transcriptional programs. **(A)** Distribution of the candidate chromosome (Chr)11 amplification (Amp.) in cancer cells per treatment condition. The average of hidden Markov model (HMM) copy-number inference values on chr11 is represented per cell on the UMAPs. **(B)** The percentage of cells by cluster and by treatment condition (average HMM value greater than 3.2) is shown. **(C-D)** UMAP combining data from 3 samples (PDOs from patient sample #3, CTL/Doxo./Taxol) colored by pseudotime value. Cells are ordered along an inferred trajectory; a black line connects all clusters. **(E-F)** EMT and CilioUp signature score levels along the inferred trajectory.

### Primary cilia promote chemoresistance of cancer cells that activate EMT programs

We next wished to determine whether primary cilia promote chemoresistance of cancer cells that activate EMT programs in our PDOs. Human TNBC PDOs are difficult to maintain in culture in the long-term which precluded the development of genetic strategies to genetically ablate primary cilia upon therapy in this model and incited us to use a pharmacological strategy. We could not detect any impact on ciliogenesis in PDOs of ciliobrevin A (CilA, Supplementary Fig. 4A), an inhibitor of the AAA+ ATPase motor dyneins reported to inhibit ciliogenesis among other cilium-independent processes in monolayers of cells in culture (*28, 39*). No other specific small molecule inhibitor of ciliogenesis has been reported.

For these various reasons, we decided to employ a microscopy-based screening assay to identify new potent small molecule inhibitors of primary ciliogenesis. The screening employed easy to grow human retinal cells (RPE1) amenable to large scale screening in monolayer culture and a library of 3271 compounds from the French National Chemical Library. We identified 12 hits in a primary screen and confirmed the capability of 9 of the drugs to reduce the number and length of primary cilia without displaying major cytotoxic effect in the cells in a secondary screen (Supplementary Fig. 4B-D). Most importantly, 5 of the drugs belonged to the same chemical family that we named Naonedin (Nao-1, Nao-2, Nao-3, Nao-4, Nao-5, Supplementary Fig. 4E). We next proceeded to determine whether one of the drugs represses primary ciliogenesis in our PDOs composed of cells that activate EMT to reside in hybrid E/M states. Nao-3 significantly repressed cilium assembly in all patient samples analyzed (Fig. 4A-B). Thus, we characterized a novel family of small-molecules that represses primary ciliogenesis both in monolayer cell culture and in patient-derived samples. We next determined whether Nao-3 suppresses chemoresistance of quasi-mesenchymal cancer ciliated cells in our PDOs. We treated PDOs with a vehicle control, Nao-3 alone, taxol alone or a combination of taxol and Nao-3. Using immunostaining for cleaved-caspase 3, we found that Nao-3 induced a modest but significant increase in the number of death associated with caspase activation of cancer cells on its own, comparable to taxol (Fig. 4C-D). Most importantly, we found that Nao-3 induced a significantly more profound death of cancer cells when combined with taxol (Fig. 4C-D). Altogether, our findings reveal a novel family of small-molecule inhibitors of primary ciliogenesis that suppresses chemoresistance of malignant cells. They offer compelling evidence that primary cilia promote chemoresistance of cancer cells that activate EMT to reside in hybrid E/M states.

**Figure 4.**
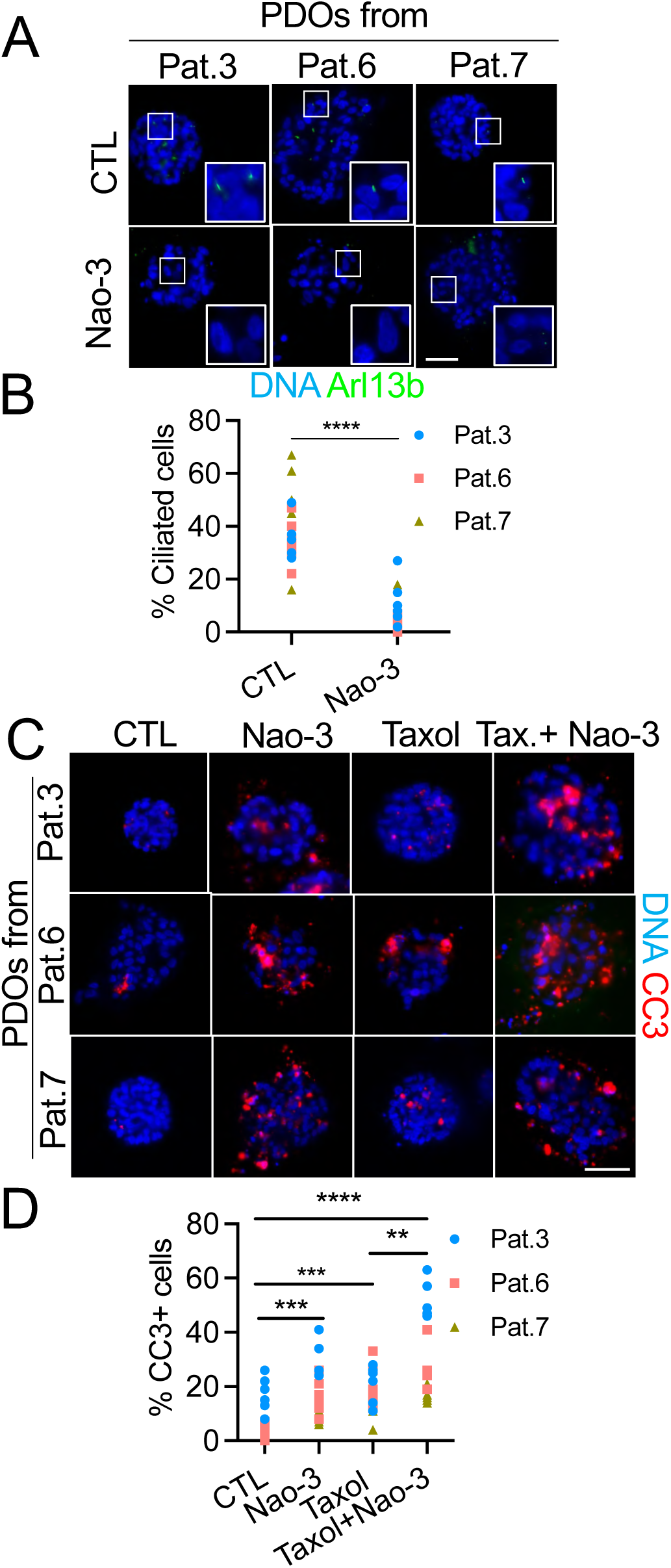
Primary cilia promote chemoresistance of hybrid E/M cancer cells. **(A)** PDOs from 3 distinct patient samples were treated with DMSO (CTL) or Naonedin-3 (Nao-3) and stained for Arl13b. Scale bar: 50 µm. **(B)** The percentage of ciliated cells was quantified. ****p ≤ 0.0001 (n=6 PDOs/patient sample/treatment condition). **(C)** PDOs were stained for the indicated proteins after the indicated treatments (Tax. = Taxol, Nao-3. = Naonedin-3). **(D)** The percentage of cleaved-caspase 3 (CC3)+ cells was measured in individual PDO for the distinct patient samples (n=6 PDOs/patient sample/treatment condition). Scale bar: 50 µm. ** p ≤ 0.01; *** p ≤ 0.001; ****p ≤ 0.0001.

### EMT-driven cell plasticity enables primary ciliogenesis in quasi-mesenchymal cells to mediate chemoresistance

To investigate whether EMT-induced primary ciliogenesis mediates chemoresistance in quasi-mesenchymal cells in complementary experiments, we next used experimentally transformed human mammary epithelial cells (HMLER). In general, the majority of HMLER cancer cells propagated *in vitro* display epithelial characteristics. In addition, we previously generated HMLER variants in which EMT can be induced experimentally by the knockdown of *Ecad* (*28*). Thus, HMLER cells acquire a quasi-mesenchymal phenotype and generate tumors that display the hallmarks of TNBC/claudin-low breast cancers when orthotopically implanted in the mouse (*27, 28*). Using epithelial (shCTL) and quasi-mesenchymal (shEcad) HMLER variants, we confirmed that while the majority of epithelial shCTL cells display epithelial characteristics, shEcad cells acquire a quasi-mesenchymal phenotype upon Ecad silencing, as judged by morphology and western blotting for epithelial and mesenchymal markers (Fig. 5A). Additionally, we demonstrated that HMLER shEcad cells gain the ability to form primary cilia using immunostaining for Arl13b (Fig. 5B).

**Figure 5.**
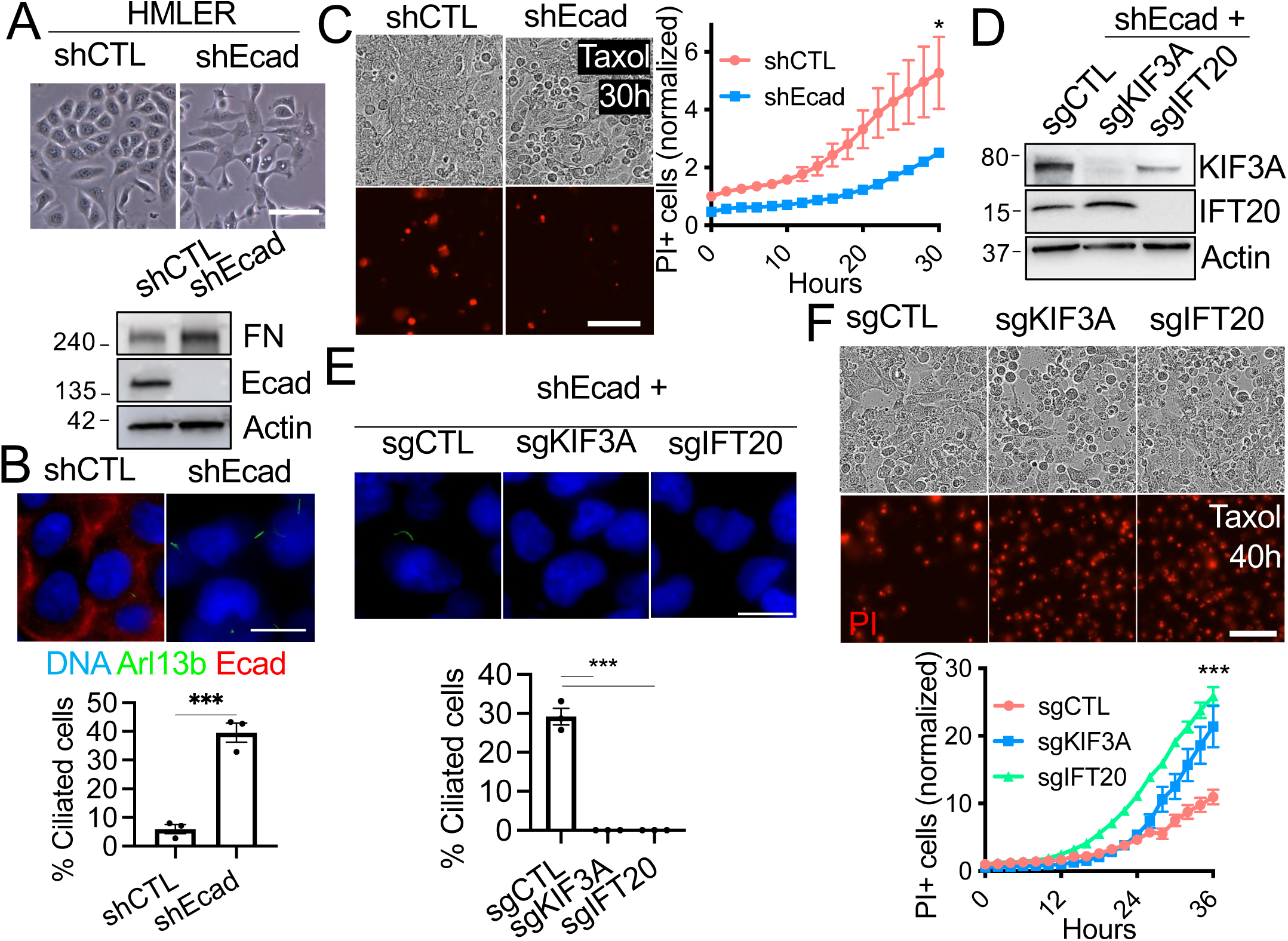
EMT-driven cell plasticity enables primary ciliogenesis in quasi-mesenchymal cells to mediate chemoresistance. **(A)** Morphology and western blot analysis of EMT markers in E-like (shCTL) and M-like (shECAD) HMLER cells. Scale bar: 100 µm. **(B)** Cells were stained for the indicated proteins to determine the percentage of ciliated cells (n=3, mean ± sem). Scale bar: 15 µm. **(C)** The sensitivity of shCTL and shEcad HMLER cells to taxol was determined by quantifying incorporation of propidium iodide (PI) in cancer cells in response to taxol (1µM) overtime (30 h) (n=3, mean ± sem). Results are normalized to the HMLER shCTL. Scale bar: 100 µm. **(D)** *KIF3A* and *IFT20* knockouts were validated by western blot of extracts for the indicated HMLER variants. **(E-F)** The impact on ciliogenesis and on taxol sensitivity was assessed as described in **(B)** and **(C)**. Scale bar: 15µm (E) and 100 µm (F). * p ≤ 0.05; *** p ≤ 0.001.

To determine whether EMT program that induces primary ciliogenesis also induce chemoresistance in HMLER cells, we next assessed the sensitivity of our epithelial shCTL versus quasi-mesenchymal ciliated shEcad HMLER variants to taxol by monitoring cell death through propidium iodide staining (Fig. 5C). We found that HMLER cells that resided in a quasi-mesenchymal state and formed primary cilia are significantly more resistant to cell death in comparison to control HMLER cells in the epithelial-like state (Fig. 5C, Supplementary Movie 2).

To investigate whether assembly of primary cilia in response to EMT is responsible for taxol resistance, we used HMLER shEcad variants in which we inhibited ciliogenesis through the knockout of *KIF3A* and *IFT20*, two genes that are essential genes for ciliogenesis. We demonstrated that knockout of these genes resulted, as anticipated, in loss of KIF3A and IFT20 proteins and ciliogenesis inhibition in quasi-mesenchymal cancer cells (Fig. 5D-E). Importantly, we found that HMLER shEcad variants in which ciliogenesis had been repressed (i.e., sgKIF3A, sgIFT20) died more of taxol relative to control ciliated HMLER shEcad cells (sgCTL, Fig. 5F, Supplementary Movie 3). Thus, primary ciliogenesis inhibition sensitize quasi-mesenchymal cancer cells to chemotherapy. Altogether, these data reveal that EMT-driven cancer cell plasticity enables primary ciliogenesis in quasi-mesenchymal cells and chemoresistance and that chemotherapy resistance is mediated by primary ciliogenesis.

## Discussion

Our previously reported data established that EMT programs induce TNBC/claudin-low tumorigenesis by inducing primary ciliogenesis in tumor-initiating cells in a murine carcinoma model (*27, 28*). Our data now emphasize a clear connection between EMT and primary ciliogenesis in human TNBCs. We demonstrate that the propensity of cancer cells to form primary cilia follows the entrance of cancer cells into a quasi-mesenchymal state (Supplementary Fig. 5). The role of the EMT/primary cilium axis in tumor response to therapy had never been addressed before. We reveal that primary cilia promote EMT-induced tumor resistance to therapy in TNBCs (Supplementary Fig. 5). Additionally, we reveal a novel family of small molecules that repress primary ciliogenesis and suppress chemoresistance. Collectively, these findings establish novel aspects of the cell biology of tumor resistance to therapy which can be targeted pharmacologically.

Our data do not exclude the possibility that our newly identified ciliogenesis inhibitor triggers the death of malignant cells through ciliogenesis inhibition and other cilium-independent functions. Nao-3 share similarities with microtubule targeting drugs carbazoles and could thus repress primary ciliogenesis by disrupting ciliary and non-ciliary microtubule dynamics (*40*). However, our work clearly establishes that primary cilia ablation on its own induces the death of malignant chemotherapy resistant cancer cells through genetic strategies. Our work is consistent with independent recent studies that reported a role for primary cilia in resistance to therapy in lung, rhabdoid and glioblastoma cancer cell lines *in vitro* (*41–43*). Jenks et al showed that in carcinoma cells of the lung, ciliogenesis is induced in response to kinase inhibitors and primary cilia mediate resistance to the drugs (*41*). The underlying reason for ciliogenesis induction in this study remained unclear but EMT-driven tumor cell heterogeneity and plasticity in response to therapy could be responsible for ciliogenesis and cilium-induced therapy resistance such as in TNBCs as demonstrated in our work. The few published studies and our data on human tumor samples provide strong evidence for the notion that primary cilia promote cancer cell resistance to various cancer therapeutics in tumors. Our data reveal new insights into mechanisms of ciliogenesis induction in response to therapy in carcinomas and reveal new pharmacological tools to repress this process which drives therapy failure. It is important to consider that targeting ciliogenesis in combination to chemotherapy and not alone appears to be to a valuable strategy given heterogeneity of cancer cells.

Targeting ciliogenesis or ciliary signaling alone could also lead to therapeutic resistance. Ciliary signaling inhibition has been used in the clinic for the treatment of basal cell carcinomas and medulloblastomas (*44, 45*). Some of these hedgehog-dependent tumors rely on primary cilia and smoothened ciliary localization for tumorigenesis (*46–49*). Inhibition of smoothened efficiently repress tumor formation but resistance can arise (*50, 51*). Interestingly, genetic screens and genomic analysis have provided evidence that loss of primary cilia due to genetic mutation in ciliary genes result in a major switch in cell signaling dependencies which can result in acquired resistance to smoothened inhibitors in these tumors (*50, 51*). Thus, combination of distinct therapies including conventional and new therapeutic strategies which target ciliogenesis/ciliary signaling and independent signaling modalities appears of great importance to successfully treat primary cilia-dependent tumors.

Pioneering work conducted in cancer cell lines *in vitro* and using genetically-engineered mouse cancer models suggested that EMT in cancer is a major route toward therapy resistance (*2, 16, 26*). Our data offer new evidence for the role of EMT in promoting inter- and intratumor phenotypic heterogeneity and therapeutic resistance in human tumors. We used PDOs to conduct our research. They recently emerged as attractive tumor models for studying tumorigenesis and therapeutic response *ex vivo* (*52–54*). Our data offer new evidence which demonstrate that PDOs represent appropriate tumor avatars for studying tumor therapeutic response. They faithfully recapitulate EMT-associated inter- and intratumor phenotypic heterogeneity found in tumors which is responsible for therapy resistance. Thus, PDOs represent useful models to reinforce our understanding on the causes and consequences of cell heterogeneity and plasticity program in human tumor and for the development of new therapeutic strategies which target EMT and related processes.

Our data offer new insights into the mechanisms of EMT-driven therapeutic resistance in human TNBCs. This group of breast neoplasms represent the most aggressive subtype of breast carcinomas which remain associated with an unmet medical need (*32*). The recent development of new therapeutics including antibody-drug conjugates enabled important progresses for the treatment of TNBCs (*31*). However, most advanced tumors remain associated with therapeutic resistance and cancer-related death (*31*). Our work emphasizes the importance of considering tumor heterogeneity and plasticity of TNBCs for therapy.

## Methods

### Human samples

Patient-derived tissues were collected from breast cancer patients that were diagnosed at the Centre Eugène Marquis and at the Institut de Cancérologie du Grand Ouest. None received therapy prior to surgery. Tissues were collected by a pathologist after resection by a surgeon. Some fragments were paraffin-embedded, other fragments were dissociated within 2 h after surgical resection (tumors). Briefly, breast tumor pieces were cut into small fragments (< 2 mm^3^), which were then dissociated enzymatically and mechanically as described previously (*27*). Viable cells were then cryopreserved before organoid culture or directly used for organoid growth.

### Patient-derived organoid culture and treatments

PDO culture was performed using a protocol adapted from Sachs and colleagues (*52*). Briefly, cells were seeded in advanced DMEM/F12 (Thermo-Fisher) supplemented with glutamax, HEPES, 10% R-spondin 1, 10% noggin, 2% B-27, nicotinamide (10 mM), N-acetyl-L-cysteine (500 mM), primocin (100 µg/mL), Y-27632 (5µM), heregulin β1 (5 nM), FGF-7 (5 ng/mL), FGF-10 (5 ng/mL), A83-01 (0.5 µM), EGF (5 ng/mL), SB202190 (1 µM) containing 5% matrigel. 200,000 cells were seeded per well in 24-well ultralow attachment plates (Corning). Organoids were analysed between passage 0 and passage 7.

### Immunofluorescence and Image Analysis

PDOs and cells were fixed and stained according to our published protocols (*27, 55, 56*). The following primary antibodies were used: Arl13b (NeuroMab 73-287, 1:200), acetylated tubulin (Cell Signaling Technology 5335; 1:1,000), E-cadherin (Cell Signaling 3195, 1:200), α-tubulin (Sigma-Aldrich T9026; 1:500), cleaved caspase 3 (Cell Signaling 9131; 1:200) and vimentin (Dako M0725; 1:500). Secondary antibodies were anti-mouse IgG2A 488 (Thermo-Fisher A21131; 1:1,000), anti-mouse 647 (Thermo-Fisher A21236, 1:1000), anti-mouse IgG1 647 (Thermo-Fisher A21240; 1:1,000) and anti-rabbit 546 (Thermo-Fisher A11035, 1:500). Mounted coverslips with cells were examined using 60X objective and a wide-field Zeiss microscope. Organoids were embedded in low-melting point agarose and analysed using a Lightsheet Z1 Zeiss microscope. Z-stacks were deconvolved and analysed with ImageJ and Imaris.

### Single-cell RNA sequencing

Single cell RNA sequencing was conducted using the Chromium Single-Cell 3’ v3.1 kit from 10X Genomics, according to the manufacturer’s protocol. Libraries were sequenced on the NovaSeq 6000 platform (Paired-end, 28 bp Read1, 90pb Read2). Raw BCL files were demultiplexing and mapped to the reference genome (refdata-cellranger-GRCh38-3.0.0) using the Cell Ranger Software Suite (v.6.1.2). The raw data were extracted from 10X format files using the *Read10X* function from the Seurat package (v4.4.0) in R v.4.1.1. We applied filtering to the feature-barcode gene expression matrix to preserve only cells of high quality based on several metrics, including the number of unique molecular identifiers (UMI), the number of genes detected and the percentage of read mapped to mitochondrial genes. Threshold were set three median absolute deviations (MAD) using the *isOutlier* function from the scater R package (v.1.22.0) to identify and flag cells with UMI counts and genes detected per cell bellow and above this specified threshold. Additionally, cells surpassing a threshold of three MAD for the percentage of mitochondrial reads were considered as non-viable and removed from the analysis. Doublets were identified using the scDblFinder R package (v.1.6.0), where the scDlbFinder score was calculated for each cell and the scDblFinder threshold was applied for doublet identification. Detailed quality control metrics are provided in Supplementary Table 6.

### Single-cell RNA sequencing data analysis

After data preprocessing, normalization was performed using the *NormalizeData* function, with default parameters, which implements a logarithmic normalization to the gene expression data. Data were centered and scaled using *ScaleData* function. During scaling step, regression of S and G2M phase scores was performed to minimized the influence of cell cycle effects in downstream analysis. For all cells from PDOs from patient sample #3 (n= 4888 cells) and cells from the three samples before (CTL n = 142 cells) and after treatment (Doxo. n= 264 cells and Taxol n= 414 cells), we used the top 2000 highly variable genes from the normalized expression matrix, to perform the principal component analysis (PCA, *runPCA* Seurat function), and applied Louvain graph-based clustering on the 30 first principal components. The resulting data were projected onto an Uniform Manifold Approximation and Projection (UMAP) using the *runUMAP* function. We applied the FindNeighbords and *FindClusters* functions with a resolution of 0.2 for the untreated PDOs from patient #3, and a resolution of 0.9 for the merged dataset of PDOs samples after treatment (CTL, Doxo, Taxol) to identify distinct clusters. For each defined cluster, a gene markers analysis was conducted using the *FindAllMarkers* function with the parameters set as follows: only.pos = TRUE, min.pct=T, logfc.threshold = 0.25. The top 5 markers genes per cluster were selected on the average log2fold change and a heatmap was generated using the *DoHeatmap* function. The score for the EMT, CilioUp and Lim_mammary_stem_cell signatures was calculating using the *AddModuleScore* function. For statistical tests comparing signature values between clusters, we utilize the *stat_compare_means* function from ggpubr R package (v.0.6.0). We employed the Wilcoxon test to compare two groups and the Kruskal-Wallis test to compare multiple groups. Statistical significance levels are indicated as follow: not significant (ns) p>0.05; * p <= 0.05; ** p<=0.01; *** p<= 0.001; **** p <= 0.0001.

Copy number variation analysis was performed using the InferCNV R package (v.1.16.0 https://github.com/broadinstitute/infercnv). The analysis was executed using the *run* function with this following parameter settings: denoise=TRUE, HMM=TRUE, leiden_resolution=1, leiden_method=”simple”, leiden_function=”modularity”, analysis_mode=”cell”. Reference cells were selected from the N-1105-epi sample, using a publicly available dataset (Pal et al., 2021) https://doi.org/10.15252/embj.2020107333

Pseudotime analysis was conducted using the R package slingshot (v.2.0.0) to perform single cell trajectory analysis, utilizing the principal curves algorithm. The analysis was conducted using the following input: cluster labels derived from the resolution of 0.9 and reduced dimensions obtained from PCA, as described previously. UMAP coordinates were used and no specific starting point was selected, default parameters were used for the analysis. ComplexHeatmap (v.2.15.4), ggplot2 (v.3.4.4), dittoSeq (v.1.4.4) R packages were used for graphical representation.

### 2D Cell Culture and treatments

HMLER cells were cultured in 1 :1 mixture ofin Dulbecco’s modified Eagle medium (DMEM/F12) supplemented with glutamax, 10% FBS, 0.01 mg/mL insulin, 0.48 µg/mL hydrocortisone, and complete mammary epithelial cell growth medium (MEGM) supplemented with bovine pituitary hormone (Lonza). For all ciliogenesis assays, cells grown until high confluence and serum starved. To repress ciliogenesis, inhibitors were added to DMEM/F12 medium without serum for 24 hours.

### Drug screening

RPE1 cells were cultured in DMEM/F12 supplemented with glutamax, 10% FBS and and penicillin/streptomycin. Cells were grown until high confluence and serum starved in DMEM/F12 for 24 hours. Drugs diluted in DMEM/F12 (10 μM) were added to the cells for 24 hours. Cells were fixed and stained as discussed above. Ciliogenesis was analyzed using a custom-made Fiji plugin.

### Viability analyses

Viability of cancer cells in PDOs was measured through a 3D CellTiter-Glo assay according to the manufacturer’s instructions. Viability of HMLER cells was assessed in 96-well flat-bottom plate (zell-kontakt). The cells were plated and then treated 24 h later with 0.1 µM and 1 µM Taxol or 0.1% DMSO in the presence of propidium iodide (Miltenyi Biotec). The plate was incubated in an IncuCyte Live Cell Analysis System (Essen Bioscience). Phase-contrast and fluorescence images of cells were acquired every 2 hours for 30-40 hours using the IncuCyte Zoom automated imaging system. Live cell analysis Incucyte software was used for data analysis.

### Western Blot Experiments

These experiments were conducted using standard procedures as described previously(*27*). Western Blot were performed using primary antibodies against KIF3A (Proteintech 13930-1-AP; 1:1,000), IFT20 (Proteintech 13615-1-AP; 1:100), E-cadherin (Cell Signaling 3195; 1:1,000), fibronectin (BD Biosciences 610077; 1:11,000) and actin (Millipore MAB1501; 1:2,000) and secondary antibodies horseradish peroxidase-coupled anti-mouse (Jackson ImmunoResearch 115-035-006; 1:5,000) or anti-rabbit (Jackson ImmunoResearch 111-035-006; 1:5,000).

### Statistical analysis

Prism was used to analyze data, draw graphs, and perform statistical analyses unless otherwise specified. Data are presented as means ± SEM. Statistical analyses were carried out by Student’s t test unless otherwise specified. *P ≤ 0.05, **P ≤ 0.01, and ***P ≤ 0.001 were considered significant. Representative results of three or more independent experiments are shown. Power analysis was conducted for all statistical studies. Blinding of sample names was used to reduce unconscious bias whenever possible.

## Acknowledgments

We thank BioCore (US16 Inserm – UAR 3556 CNRS) and Biosit (US18 Inserm - UAR 3480 CNRS) biotechnology centers, including MicroPICell, MRic, H2P2, CRB Santé core facilities, as well as GenoA the genomics core facility for technical support. This work was supported by Fondation de France, Région Pays de la Loire, SIRIC ILIAD INCa-DGOS-INSERM-ITMO Cancer_18011, Inserm, Ligue Contre le Cancer, Institut de Cancérologie de l’Ouest, Cancéropôle Grand Ouest.

## Author contributions

Conceptualization, C.E.T, J.D., C.P., R.A.W., J.A.L., P.P.J., V.J.G.; Methodology, C.E.T, J.D., C.E.H, X.P., M.L.G., F.G., P.T., C.P., M.C., R.A.W., J.A.L., P.P.J., V.J.G.; Investigation, C.E.T., J.D., A.M.M.D., M.M, J.R., E.D., C.E.H., X.P., R.V., P.S., V.J.G.; Writing – Original Draft, C.E.T., J.D., V.J.G., Review & Editing, XX; Funding Acquisition, C.P., M.C., J.A.L., P.P.J., V.J.G.; Resources, M.L.G., F.G., P.T., P.S., P.D., M.C., R.A.W.; Supervision, V.J.G.

---

**Supplementary Fig. 1.**
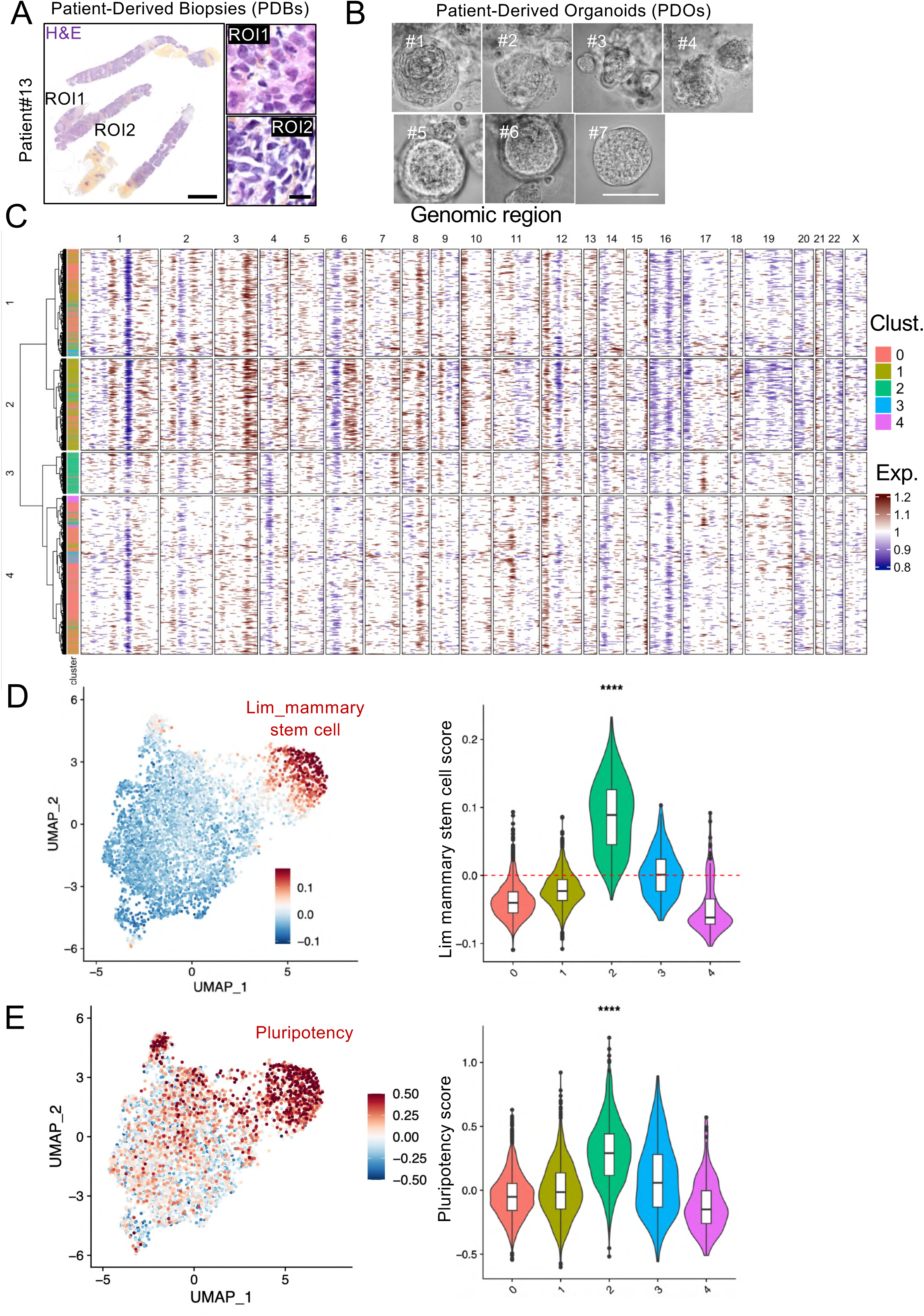
Cancerous lesions in patient-derived biopsies and additional features of patient-derived cancer organoids. **A)** Paraffin sections from patient-derived biopsies (PDBs) were stained with H&E (n = 13, a representative result for patient#13 is shown), Scale bars : 2 mm (low magnification) and 100 μm (high magnification). **(B)** The morphology of PDOs from distinct patient samples (#1-7) was examined by brightfield microscopy. Scale bar: 100 μm. **(C)** Heatmap showing inferred CNVs in cells of PDOs from patient sample#3 with unsupervised hierarchical clustering. **(D-E)** UMAP and violin plots illustrating the expression of mammary stem cell and pluripotency signatures in the distinct cell clusters composing PDOs from patient sample#3.

**Supplementary Fig. 2.**
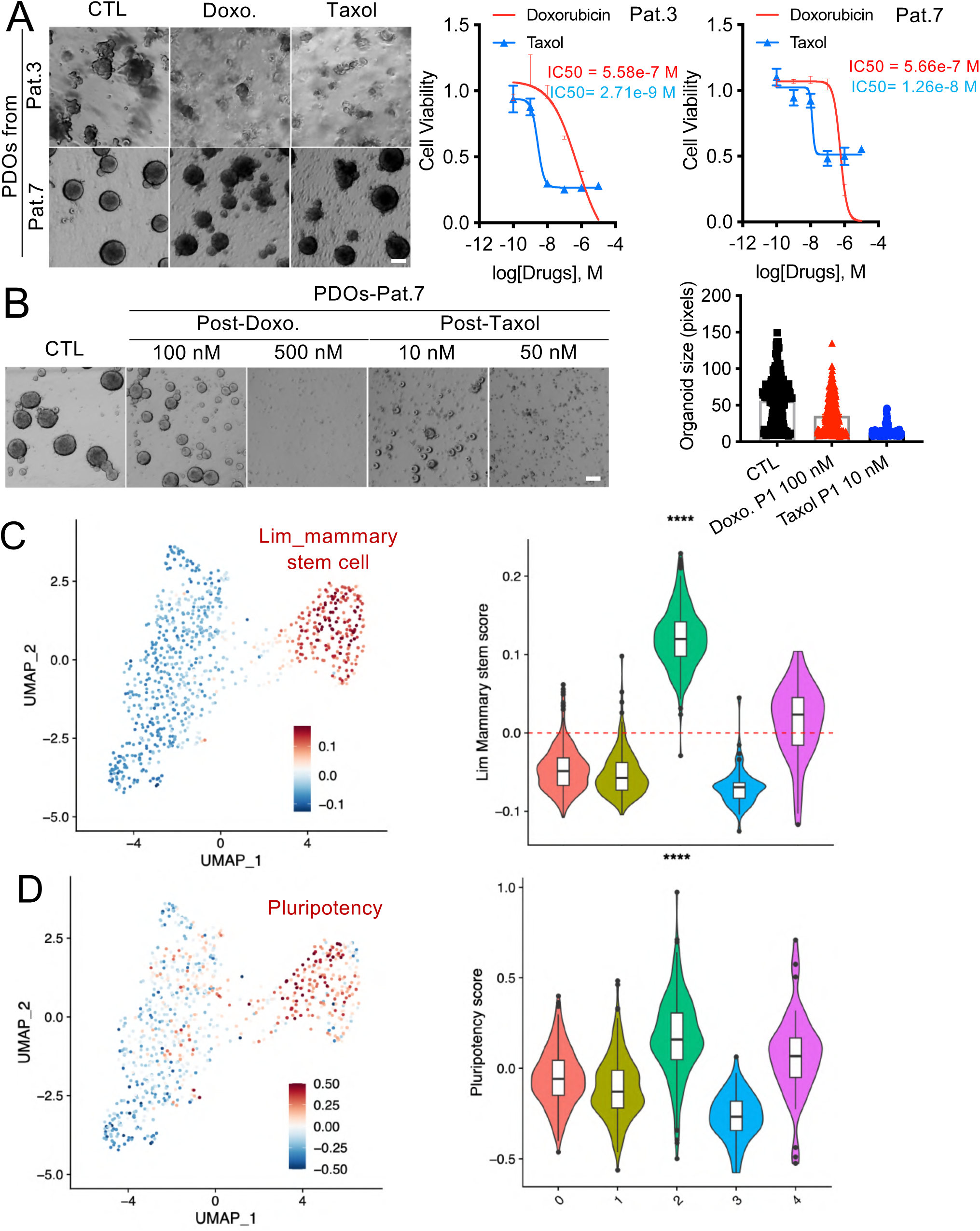
Impact of chemotherapy on PDOs and genomic features of quasi-mesenchymal chemoresistant cells. **(A)** The dose-dependent impact of chemotherapy on cell viability in PDOs was tested for two distinct patient (pat.) samples (#3 and #7). Representative images of PDOs treated with DMSO (CTL) or with doxorubicin (doxo., 100 nM) and taxol (10 nM) after 48h of treatment are shown. The cell viability was measured at different concentrations and the IC50 is shown for each drug (mean ± sem, representative analysis of 3 independent experiments). Scale bar: 100 μm. **(B)** The impact of drugs on the ability of chemoresistant cancer cells to reconstitute PDOs after drug removal at two intermediate drug concentrations is shown and measured at the indicated concentrations. Scale bar: 100 μm (n ≥ 213 PDOs/treatment condition, mean ± sem). **(C-D)** UMAP and violin plots illustrating the expression of mammary stem cell and pluripotency signatures in the distinct cell clusters from PDOs post-treatment.

**Supplementary Fig. 3.**
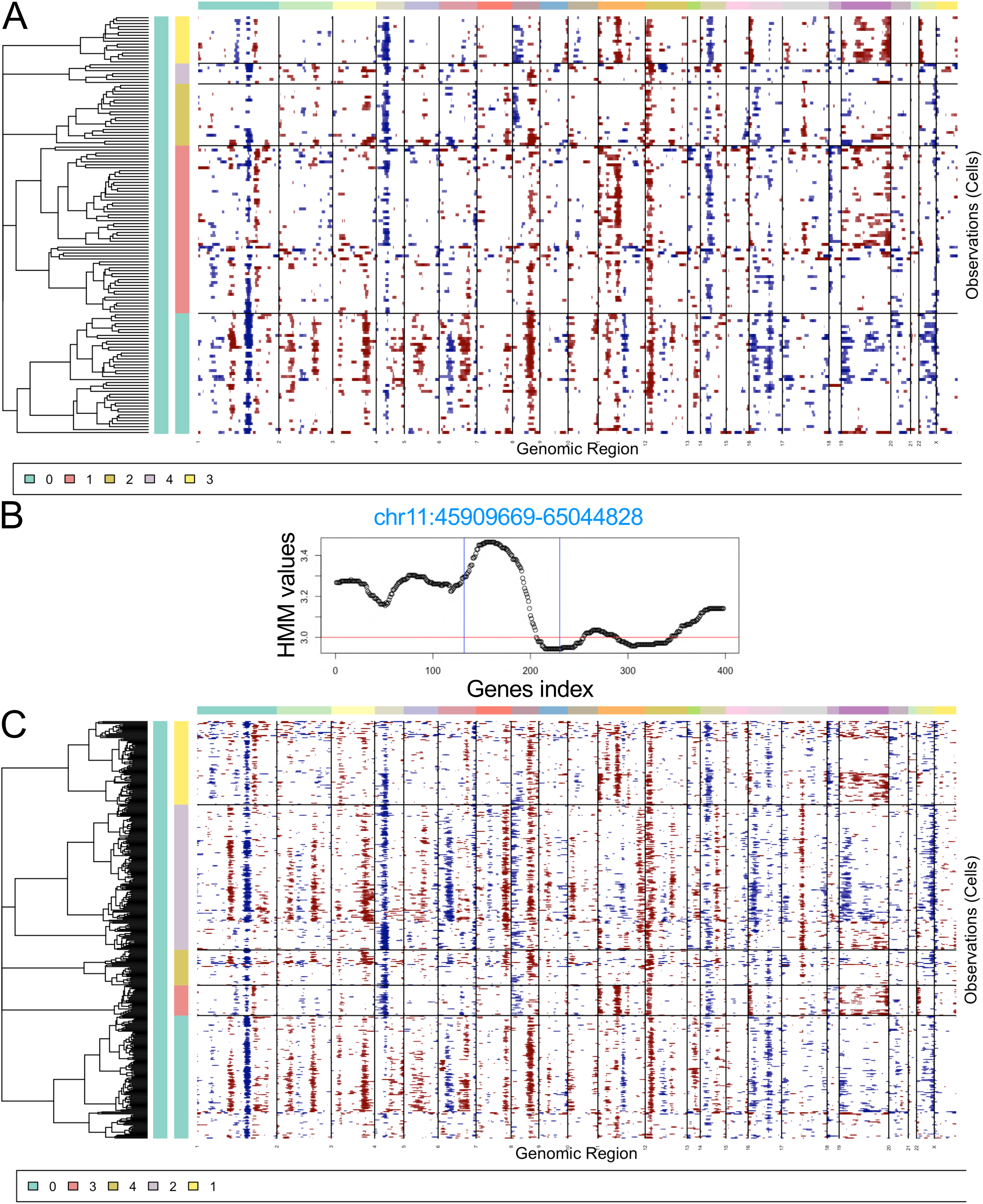
Analysis of copy number aberrations inferred upon chemotherapy. **(A)** Heatmap displaying inferred CNVs in cells of PDOs from the control sample categorized by cluster. **(B)** Average values of HMM copy number inference on chromosome 11 for all cells in the control sample. The x-axis represents the order of genes. **(C)** Heatmap showing inferred CNVs in cells of CTL and chemotherapy treated PDOs categorized by cluster.

**Supplementary Fig. 4.**
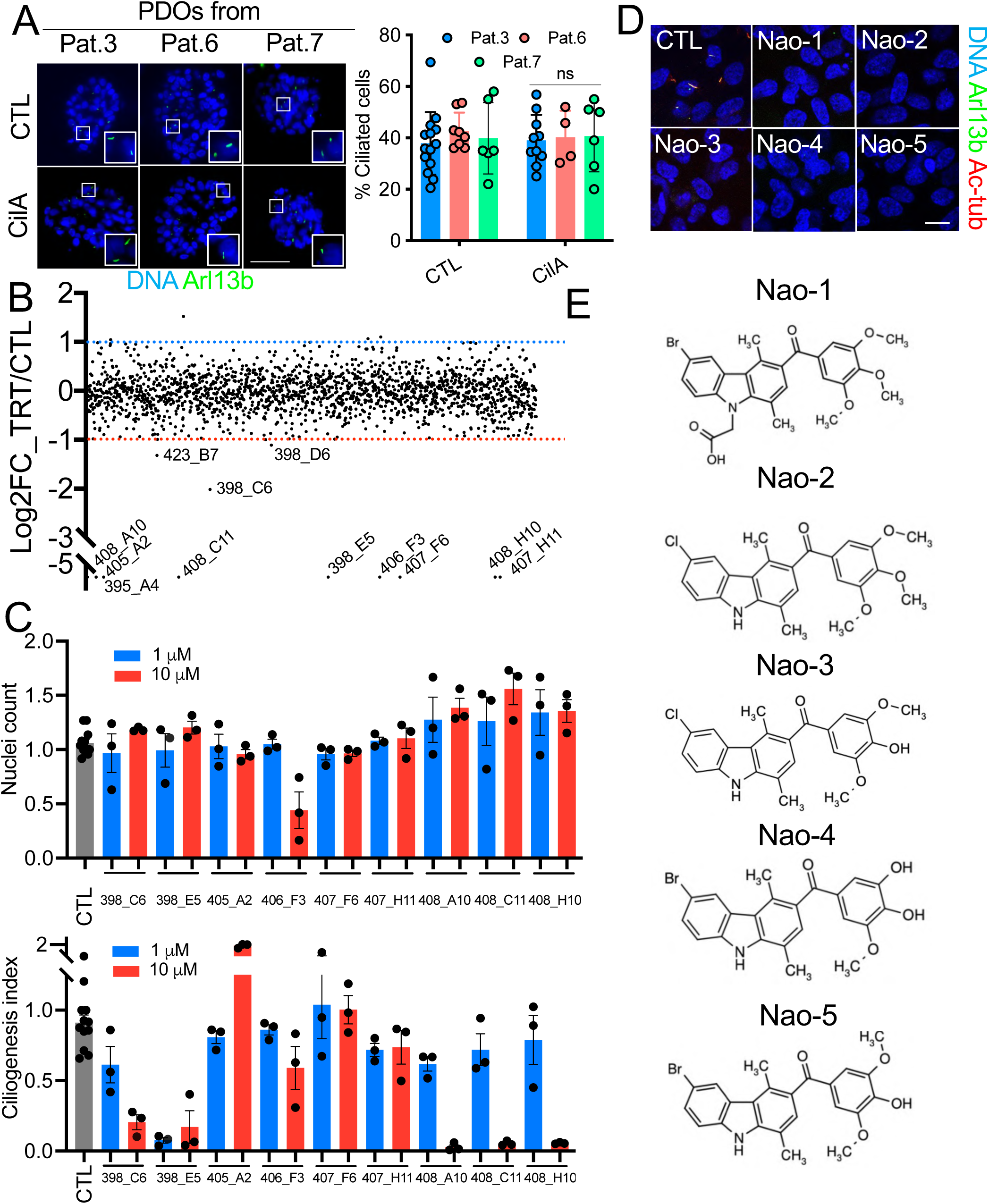
Identification of small-molecule inhibitors of primary ciliogenesis. **(A)** PDOs from distinct patient (pat.) samples were treated with DMSO (CTL) or Ciliobrevin A (CilA, 100 µM, 24h) and stained for Arl13b. Scale bar: 50 µm. The percentage of ciliated cells was quantified in distinct PDOs for each patient sample. ns p > 0.05 (n ≥ 4 PDOs/patient sample/treatment condition, means ± sem). **(B)** RPE1 cells were plated at high confluence. They were serum starved 24h post-plating to induce ciliogenesis. Cells were treated 24h post-starvation with 3271 chemical compounds (10 µM). They were fixed and stained 24h post-treatment for DNA, Arl13b and acetylated-tubulin. Nuclei number and the percentage of ciliated cells was determined based on Arl13b staining using a custom-made Fiji plugin. Compounds which markedly repressed ciliogenesis (Log2 fold change (FC) in the percentage of ciliated cells in treated (TRT) versus Control (CTL, DMSO treated) cells are indicated below the red line. **(C-D)** A secondary screen with the hits from the primary screen at two distinct concentrations (1 and 10 µM) was conducted (as described in B). The number of cells as well as the ciliogenesis index (computational analysis of the percentage of ciliated cells) was determined 24h after each treatment. Results are relative to CTL, DMSO-treated cells (mean ± sem). The impact of the most potent inhibitors on ciliogenesis is shown at the indicated concentration. Scale bar: 15 µm. **(E)** Chemical structure of Naonedins.

**Supplementary Fig. 5.**
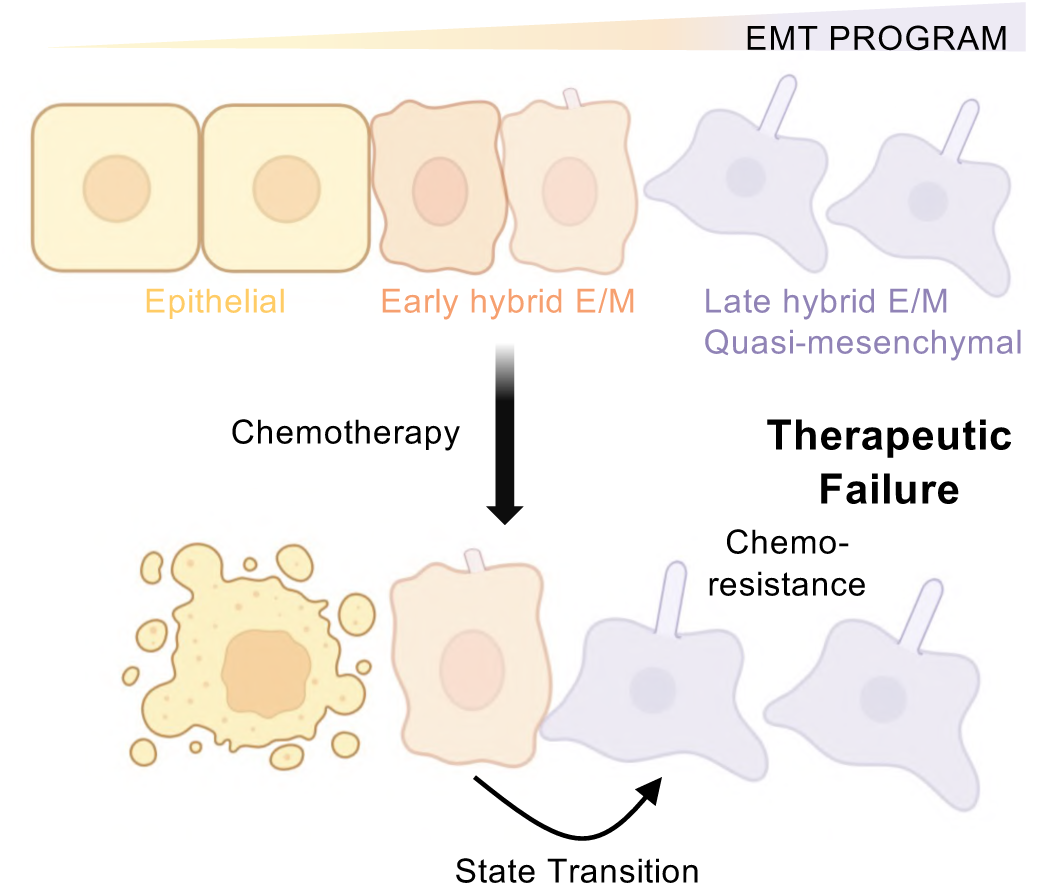
Schematic representation of phenotypic plasticity of cancer cells mediating therapeutic resistance in human TNBC. Our results show that in cancer cells of human TNBC, activation of EMT enables malignant cells to acquire distinct hybrid E/M phenotypes. Late-hybrid quasi-mesenchymal cells assemble primary cilia. Chemotherapy can induce cell state transition from the early hybrid to the late hybrid E/M phenotype. Primary cilia promote chemoresistance of cells that reside in a late hybrid quasi-mesenchymal state and help TNBC evade therapy.

## References

1. H. Bonnefoi et al., Pathological complete response after neoadjuvant chemotherapy is an independent predictive factor irrespective of simplified breast cancer intrinsic subtypes: a landmark and two-step approach analyses from the EORTC 10994/BIG 1-00 phase III trial. Ann Oncol 25, 1128–1136 (2014).

2. J. C. Marine, S. J. Dawson, M. A. Dawson, Non-genetic mechanisms of therapeutic resistance in cancer. Nat Rev Cancer 20, 743–756 (2020).

3. F. Bray, M. Laversanne, E. Weiderpass, I. Soerjomataram, The ever-increasing importance of cancer as a leading cause of premature death worldwide. Cancer 127, 3029–3030 (2021).

4. I. Dagogo-Jack, A. T. Shaw, Tumour heterogeneity and resistance to cancer therapies. Nat Rev Clin Oncol 15, 81–94 (2018).

5. C. Kim et al., Chemoresistance Evolution in Triple-Negative Breast Cancer Delineated by Single-Cell Sequencing. Cell 173, 879–893 e813 (2018).

6. B. B. Liau et al., Adaptive Chromatin Remodeling Drives Glioblastoma Stem Cell Plasticity and Drug Tolerance. Cell Stem Cell 20, 233–246 e237 (2017).

7. J. Marsolier et al., H3K27me3 conditions chemotolerance in triple-negative breast cancer. Nat Genet 54, 459–468 (2022).

8. I. Pastushenko et al., Identification of the tumour transition states occurring during EMT. Nature 556, 463–468 (2018).

9. B. Bierie et al., Integrin-beta4 identifies cancer stem cell-enriched populations of partially mesenchymal carcinoma cells. Proc Natl Acad Sci U S A 114, E2337–E2346 (2017).

10. I. Pastushenko, C. Blanpain, EMT Transition States during Tumor Progression and Metastasis. Trends Cell Biol 29, 212–226 (2019).

11. C. E. Tessier et al., EMT and primary ciliogenesis: For better or worse in sickness and in health. Genesis, e23568 (2023).

12. A. Dongre, R. A. Weinberg, New insights into the mechanisms of epithelial-mesenchymal transition and implications for cancer. Nat Rev Mol Cell Biol 20, 69–84 (2019).

13. M. A. Nieto, R. Y. Huang, R. A. Jackson, J. P. Thiery, Emt: 2016. Cell 166, 21–45 (2016).

14. M. M. Wilson, R. A. Weinberg, J. A. Lees, V. J. Guen, Emerging Mechanisms by which EMT Programs Control Stemness. Trends Cancer 6, 775–780 (2020).

15. C. Kroger et al., Acquisition of a hybrid E/M state is essential for tumorigenicity of basal breast cancer cells. Proc Natl Acad Sci U S A 116, 7353–7362 (2019).

16. P. B. Gupta, I. Pastushenko, A. Skibinski, C. Blanpain, C. Kuperwasser, Phenotypic Plasticity: Driver of Cancer Initiation, Progression, and Therapy Resistance. Cell Stem Cell 24, 65–78 (2019).

17. R. M. Pommier et al., Comprehensive characterization of claudin-low breast tumors reflects the impact of the cell-of-origin on cancer evolution. Nat Commun 11, 3431 (2020).

18. M. Saxena, M. A. Stephens, H. Pathak, A. Rangarajan, Transcription factors that mediate epithelial-mesenchymal transition lead to multidrug resistance by upregulating ABC transporters. Cell Death Dis 2, e179 (2011).

19. C. A. Del Vecchio et al., De-differentiation confers multidrug resistance via noncanonical PERK-Nrf2 signaling. PLoS Biol 12, e1001945 (2014).

20. D. W. Wu et al., FHIT loss confers cisplatin resistance in lung cancer via the AKT/NF-kappaB/Slug-mediated PUMA reduction. Oncogene 34, 2505–2515 (2015).

21. M. Debaugnies et al., RHOJ controls EMT-associated resistance to chemotherapy. Nature 616, 168–175 (2023).

22. Z. Zhang et al., Activation of the AXL kinase causes resistance to EGFR-targeted therapy in lung cancer. Nat Genet 44, 852–860 (2012).

23. W. L. Tam et al., Protein kinase C alpha is a central signaling node and therapeutic target for breast cancer stem cells. Cancer Cell 24, 347–364 (2013).

24. A. Dongre et al., Epithelial-to-Mesenchymal Transition Contributes to Immunosuppression in Breast Carcinomas. Cancer Res 77, 3982–3989 (2017).

25. A. Dongre et al., Direct and Indirect Regulators of Epithelial-Mesenchymal Transition-Mediated Immunosuppression in Breast Carcinomas. Cancer Discov 11, 1286–1305 (2021).

26. T. Shibue, R. A. Weinberg, EMT, CSCs, and drug resistance: the mechanistic link and clinical implications. Nat Rev Clin Oncol 14, 611–629 (2017).

27. M. M. Wilson et al., An EMT-primary cilium-GLIS2 signaling axis regulates mammogenesis and claudin-low breast tumorigenesis. Sci Adv 7, eabf6063 (2021).

28. V. J. Guen et al., EMT programs promote basal mammary stem cell and tumor-initiating cell stemness by inducing primary ciliogenesis and Hedgehog signaling. Proc Natl Acad Sci U S A, (2017).

29. V. J. Guen, C. Prigent, Targeting Primary Ciliogenesis with Small-Molecule Inhibitors. Cell Chem Biol 27, 1224–1228 (2020).

30. K. I. Hilgendorf, B. R. Myers, J. F. Reiter, Emerging mechanistic understanding of cilia function in cellular signalling. Nat Rev Mol Cell Biol, (2024).

31. A. Bardia et al., Sacituzumab Govitecan in Metastatic Triple-Negative Breast Cancer. N Engl J Med 384, 1529–1541 (2021).

32. T. Grinda et al., Evolution of overall survival and receipt of new therapies by subtype among 20 446 metastatic breast cancer patients in the 2008-2017 ESME cohort. ESMO Open 6, 100114 (2021).

33. J. Knezevic et al., Expression of miR-200c in claudin-low breast cancer alters stem cell functionality, enhances chemosensitivity and reduces metastatic potential. Oncogene 34, 5997–6006 (2015).

34. E. Nolan, G. J. Lindeman, J. E. Visvader, Deciphering breast cancer: from biology to the clinic. Cell 186, 1708–1728 (2023).

35. J. I. Herschkowitz et al., Identification of conserved gene expression features between murine mammary carcinoma models and human breast tumors. Genome Biol 8, R76 (2007).

36. A. Prat et al., Phenotypic and molecular characterization of the claudin-low intrinsic subtype of breast cancer. Breast Cancer Res 12, R68 (2010).

37. C. Fougner, H. Bergholtz, J. H. Norum, T. Sorlie, Re-definition of claudin-low as a breast cancer phenotype. Nat Commun 11, 1787 (2020).

38. S. Z. Wu et al., A single-cell and spatially resolved atlas of human breast cancers. Nat Genet 53, 1334–1347 (2021).

39. A. J. Firestone et al., Small-molecule inhibitors of the AAA+ ATPase motor cytoplasmic dynein. Nature 484, 125–129 (2012).

40. L. Peronne et al., Two Antagonistic Microtubule Targeting Drugs Act Synergistically to Kill Cancer Cells. Cancers (Basel) 12, (2020).

41. A. D. Jenks et al., Primary Cilia Mediate Diverse Kinase Inhibitor Resistance Mechanisms in Cancer. Cell Rep 23, 3042–3055 (2018).

42. S. O. Kim, B. Y. Kim, K. H. Lee, Synergistic effect of anticancer drug resistance and Wnt3a on primary ciliogenesis in A549 cell-derived anticancer drug-resistant subcell lines. Biochem Biophys Res Commun 635, 1–11 (2022).

43. L. Wei et al., Inhibition of Ciliogenesis Enhances the Cellular Sensitivity to Temozolomide and Ionizing Radiation in Human Glioblastoma Cells. Biomed Environ Sci 35, 419–436 (2022).

44. S. J. Scales, F. J. de Sauvage, Mechanisms of Hedgehog pathway activation in cancer and implications for therapy. Trends Pharmacol Sci 30, 303–312 (2009).

45. N. Basset-Seguin, H. J. Sharpe, F. J. de Sauvage, Efficacy of Hedgehog pathway inhibitors in Basal cell carcinoma. Mol Cancer Ther 14, 633–641 (2015).

46. S. E. Conduit et al., A compartmentalized phosphoinositide signaling axis at cilia is regulated by INPP5E to maintain cilia and promote Sonic Hedgehog medulloblastoma. Oncogene, (2017).

47. Y. G. Han et al., Dual and opposing roles of primary cilia in medulloblastoma development. Nat Med 15, 1062–1065 (2009).

48. N. Yang et al., INTU is essential for oncogenic Hh signaling through regulating primary cilia formation in basal cell carcinoma. Oncogene, (2017).

49. S. Y. Wong et al., Primary cilia can both mediate and suppress Hedgehog pathway-dependent tumorigenesis. Nat Med 15, 1055–1061 (2009).

50. F. Kuonen et al., Loss of Primary Cilia Drives Switching from Hedgehog to Ras/MAPK Pathway in Resistant Basal Cell Carcinoma. J Invest Dermatol 139, 1439–1448 (2019).

51. X. Zhao et al., A Transposon Screen Identifies Loss of Primary Cilia as a Mechanism of Resistance to SMO Inhibitors. Cancer Discov 7, 1436–1449 (2017).

52. N. Sachs et al., A Living Biobank of Breast Cancer Organoids Captures Disease Heterogeneity. Cell 172, 373–386 e310 (2018).

53. J. F. Dekkers et al., Long-term culture, genetic manipulation and xenotransplantation of human normal and breast cancer organoids. Nat Protoc 16, 1936–1965 (2021).

54. S. Bhatia et al., Patient-Derived Triple-Negative Breast Cancer Organoids Provide Robust Model Systems That Recapitulate Tumor Intrinsic Characteristics. Cancer Res 82, 1174–1192 (2022).

55. M. Duclos, C. Prigent, R. Le Borgne, V. J. Guen, Three-Dimensional Imaging of Organoids to Study Primary Ciliogenesis During ex vivo Organogenesis. J Vis Exp, (2021).

56. A. M. M. Dupuy, P. P. Juin, V. J. Guen. (2022).

